# Deep learning does not outperform classical machine learning for cell-type annotation

**DOI:** 10.1101/653907

**Authors:** Niklas D. Köhler, Maren Büttner, Niry Andriamanga, Fabian J. Theis

**Author notes:** these authors contributed equally (Niklas D. Köhler, Maren Büttner).

## Abstract

Deep learning has revolutionized image analysis and natural language processing with remarkable accuracies in prediction tasks, such as image labeling and semantic segmentation or named-entity recognition and semantic role labeling. Specifically, the combination of algorithmic and hardware advances with the appearance of large and well-labeled datasets has led up to seminal contributions in these fields.

The emergence of large amounts of data from single-cell RNA-seq and the recent global effort to chart all cell types in the Human Cell Atlas has attracted an interest in deep-learning applications. However, all current approaches are unsupervised, *i.e.*, learning of latent spaces without using any cell labels, even though supervised learning approaches are often more powerful in feature learning and the most popular approach in the current AI revolution by far. Here, we ask why this is the case. In particular we ask whether supervised deep learning can be used for cell annotation, *i.e.* to predict cell-type labels from single-cell gene expression profiles. After evaluating 10 classification methods across 14 datasets, we notably find that deep learning does not outperform classical machine-learning methods in the task. Thus, cell-type prediction based on gene-signature derived cell-type labels is potentially too simplistic a task for complex non-linear methods, which demands better labels of functional single-cell readouts.

## Main

Single-cell genomics is a success story in global scientific collaborative activities and research, combining advances in genomics, miniaturization, and data science. The widespread availability of transcriptome readouts from various cells, tissues, and species nowadays allows the integration and comparison of such data using machine-learning methods^1–5^. Successful data integration in turn facilitates reliable annotation of cell types at new datasets levels of quality as compared with those of human experts. Annotating cell types is a classical supervised learning problem, which can be addressed by using numerous classification models, such as logistic regression, support vector machines, and random forests. However, with increasing data volumes, we can soon approach data volume thresholds where deep learning models outperform classical machine-learning models in cell-type annotation tasks. Here we explore whether we have reached this threshold by comparing classical machine-learning approaches with state-of-the-art deep-learning architectures in a cell-type annotation task. In our comparison study of classifier types, we present a strategy for deriving optimal parameters for a given model to predict cell-type annotation, *i.e.*, what level of accuracy can we expect in the best case scenario from a given model trained using a specific dataset. We applied this testing framework to three different biological systems with increasing sample sizes: mouse embryonic development^6–10^ (600 cells), human pancreas^11–14^ (approx. 20,000 cells), and mouse brain^15–18^ (approx. 1,000,000 cells), each with varied complexity.

### A unified strategy for cell type annotation

We consider the following general problem setup: Given the access to a set of *n*-*1* reference studies with labeled cell types, our objective is to assign unknown cells to cell types automatically in a new target study of interest. Notably, recent studies for cell type annotation tasks did not split by dataset for evaluating the prediction accuracy, therefore limiting the informative power ^19–21^. Predicting a previously unseen dataset is a classical supervised learning problem with the added complexity that, depending on the study, we may expect novel cell types (labels). We address the issue in a manner similar to scmap^1^ as follows. First, we define a confidence threshold in our prediction (*thresh* = *0.5*). Please note that every cell-type class prediction has a specific confidence level. Subsequently, we assign cells below the threshold with the tag “unknown cell type”, which indicates that the cell has minimal similarity to all the annotated cell types in the reference studies, and is therefore, potentially a novel cell type.

We analyzed five machine-learning models of increasing complexity for our comparison: k-NN classification (knn), logistic regression (lgr) (with both Lasso and ridge penalty), Support Vector Machines (svm), Gradient Boosted Trees (xgb), and Multilayer Perceptrons (mlp) (see **Supplementary Methods** for details). To ensure a meaningful and unbiased comparison, we perform a five-fold cross-validated grid search over a wide range of potential parameters for all classifiers (see **Supplementary Fig. 1**). The optimal parameter set for each classifier is selected based on the highest mean accuracy score averaged over classes and over the respective five-fold cross-validation test set predictions, therefore accounting for class imbalance. Notably, using class weighting instead did not improve the average accuracy nor the overall accuracy (see **Supplementary Table 1**). We perform the grid search approach in a leave-one-study-out fashion for all three biological systems, resulting in 14 five-fold cross-validated grid searches for five different classifier types. The classifiers with the final parameter sets are then used to predict the unknown cell types in the target study.

We further included five single-cell specific cell type annotation models: scmap^1^, SingleR^22^, Garnett^5^, scMatch^23^ and cellFishing.jl^24^. While scmap, SingleR and cellFishing.jl can be trained on *n*-*1* datasets and applied to the *n*th dataset, scMatch uses a FANTOM5 database derived reference and Garnett trains a classifier on marker genes.

### Most machine learning methods are well-suited for cell type annotation given optimal hyper-parameter values

We evaluated cell-type annotation performance of models based on the accuracy of predicting cell types in a holdout study. In the case of scMatch, we considered a cell as correctly classified, if it was assigned to the correct tissue (e.g. embryonic tissue in the case of embryonic development and pancreas for the human pancreas scenario). For the machine-learning models, we particularly measured the accuracy based on receiver operator characteristics per cell type to determine the best performing model across all classifier types (see **Fig. 1a**). Subsequently, we determined the recall of the best performing model and present it as a confusion matrix of actual and predicted cell types in the target study (see **Fig. 1b** and **Supplementary Figure 2**). The final evaluation score is the accuracy of the top performing classifier for predicting all cell-type classes calculated on the target study (see **Fig. 1c**).

**Figure 1:**
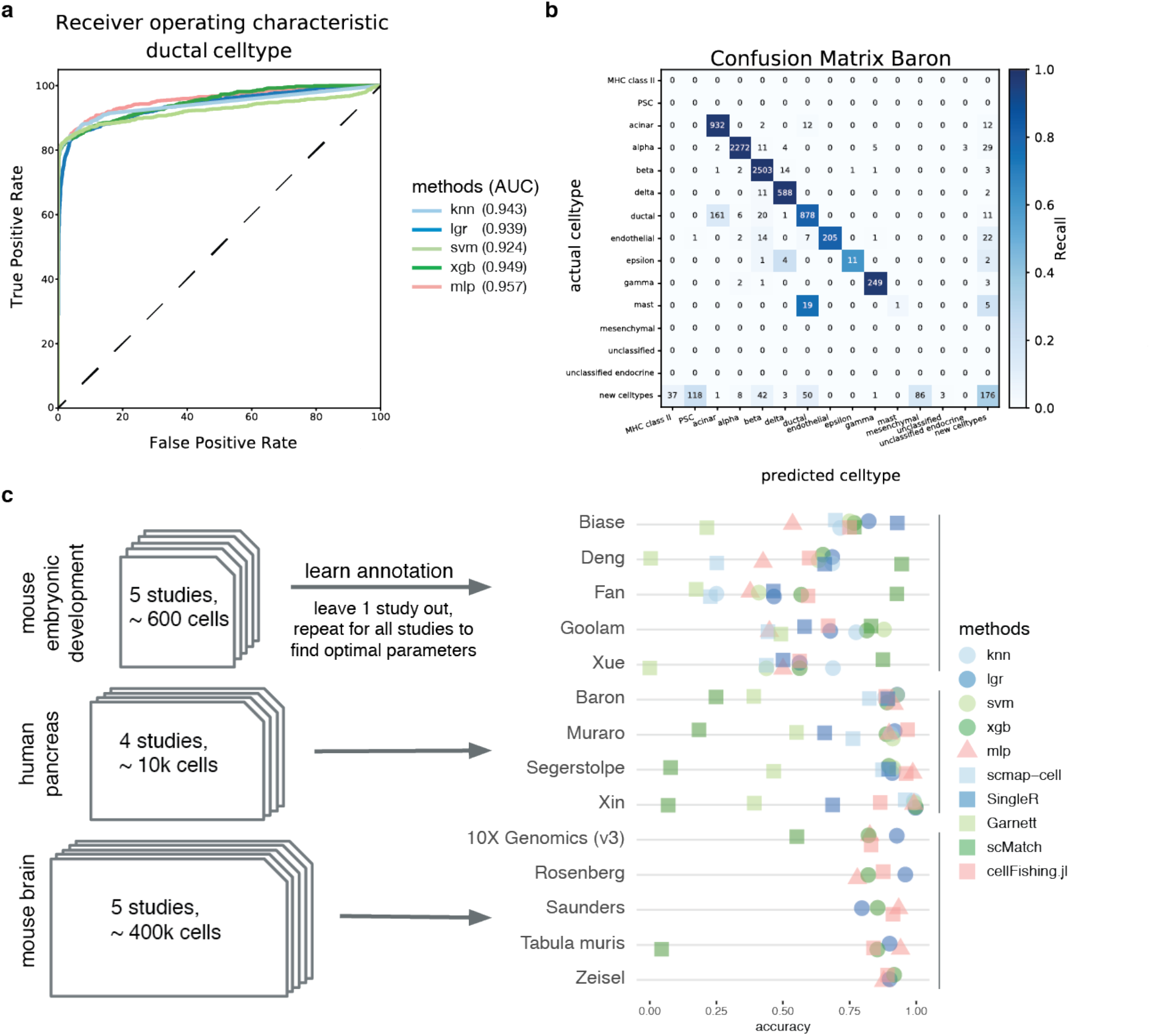
All supervised learning models for cell-type annotation have similar performance as measured by accuracy. **(a)** Receiver operating characteristics for the ductal cell type (Baron data as holdout study) in all models, which return prediction probability for all classes. The area under curve (AUC) is given in parentheses in the legend. We observe little difference across classifier types. **(b)** Confusion matrix for the recall of the MLP in pancreas data with Baron12 data as holdout study. **(c)** Accuracy scores for knn, lgr, svm, xgb (circles) and mlp (red triangles) in each study using the optimal parameter set and single-cell specific models (squares, scmap-cell, SingleR, Garnett, scMatch and cellFishing.jl). In the case of mouse brain data, only included lgr, xgb, mlp, cellFishing.jl and scMatch (2 data sets) due to infeasible runtime or memory usage.

For the small-scale embryonic development datasets, we observed considerable differences in performance across models in the same dataset, which could be explained by the small sample size of the training data. For example, the dataset from Deng et al.^8^ comprises 314 cells, and the training dataset comprises 284 cells. However, the overall accuracy of the prediction across datasets was not correlated with the sample size ratio of the training and test dataset. Instead, the differences in performance across models on the same test dataset decreased with increasing sample size in the training dataset.

When we used tissue studies of the human pancreas^11–14^ and the mouse brain^15–18^, all machine-learning models and *cellFishing.jl* performed equally well on the prediction task. Smith et al.^25^ reported a similar result in bulk RNA-seq data. In contrast, the single-cell specific classifiers *scMatch* and *Garnett* showed lower accuracy on the same datasets. We achieved a high prediction accuracy and recall in human pancreas studies. We have illustrated the cell-type prediction for the Baron^12^ dataset with MLP (see **Fig. 1b**). Notably, MLP could not predict several rare cell types at all (e.g. mast cells), while it predicted abundant cell types accurately. Notably the reference dataset was more complex and contained more cell types compared to the predicted dataset. The prediction favored rare cell types, such as mesenchymal cells and PSCs, which the authors of the original publication did not observe. Whether the prediction reflects the ground truth better than the human expert labels, is debatable. In general, reproducibility across human experts is limited ^26^ and algorithmic discovery of rare cell types remains challenging ^27^.

In case of the large mouse brain datasets, we included lgr, xgb, mlp, *cellFishing.jl* and *scMatch* (2 data sets) due to limitations in runtime and memory usage of all other models. The accuracy scores for each classifier varied minimally for every holdout study and were generally comparable across all holdout studies. We even observed a good performance on the 10X Genomics prenatal mouse brain dataset (E18 stage) even though we expected differences caused by maturation from the training data, which were obtained from postnatal mice. The developing brain undergoes considerable maturation processes before and after birth, which are reflected in the transcriptomes of the respective cells. Surprisingly, the lower transcriptional similarity between test data and training data did not result in lower prediction accuracy scores.

Overall, we found that the deep learning models did not outperform classical machine-learning models, such as logistic regression and svm (pairwise wilcoxon rank sum test, see **Supplementary Tables 2-4**), in the prediction/annotation of cell types in new data sets. Instead, factors, such as similarities between training and test data, data complexity in terms of cell types, and mere sample size affect the accuracy of annotation. Nevertheless, model tuning through grid search considerably improved accuracy. Almost all tested classifiers significantly outperformed *scmap-cell^1^* and *Garnett^5^* based on accuracy scores (pairwise wilcoxon rank sum test, see **Supplementary Tables 2-4**, see **Fig. 1c**). Interestingly, cellFishing.jl^24^ performed equally well as the general classifiers, significantly better than *Garnett* (see **Supplementary Table 2**), and was the only single-cell specific tool that scaled to all datasets in this study.

### Discussion and conclusion

We tested several supervised classifiers in small, medium, and large-scale datasets. We compared optimally tuned classification models with various degrees of complexity using a global hyperparameter search to obtain optimal parameter sets. It is not surprising that the performance of all the trained models improved significantly compared to the model *scmap-cell*, which uses a fixed set of parameters. Clearly, cell-type annotation accuracy benefits from the screening for optimal parameters independent of the model. In addition, the accuracy of the various classifiers is largely similar for each holdout study across numerous data sets, where the differences in the large datasets are lower compared to the differences in small datasets. Interestingly, the feature-based classifier Garnett and the database approach scMatch were significantly outperformed by supervised classification with cell type labels.

Why does deep learning not outperform classical machine-learning methods, in particular for the large and complex droplet-based data sets with more than half a million cells? Firstly, our input data is tabular rather than sequential. Additional information on the role of genes, as encoded in gene regulatory networks, could be leveraged e.g. in graph convolutional networks. Secondly, we have to consider when a learning task is ‘complex’ enough for powerful representation learning algorithms like deep learning to outperform simpler methods like kNN, logistic regression and SVMs. The difference between a simple and a complex learning task lies in the complexity of the transformations of the raw data necessary to identify a representation, which linearly separates the target labels of interest. Therefore, a learning task can be considered ‘complex’ when its label-separating representation has very little resemblance with the similarity structure present in the raw data. For example, learning on RGB image datasets is simple when we want to predict the amount of red vs green pixels, but a learning task on the same image dataset becomes incredibly complex when we aim to predict the content on an image. Deep learning is a very powerful representation learning technique, which excels exactly in constructing such representations: Given enough data, deep learning based methods are able to discover features, which span vastly different data graphs compared to raw data. We illustrated this well-known feature on the often-studied cifar10 dataset (60,000 images in 10 mutually exclusive classes; see **Supplementary Fig. 3**). Here, separating the classes based on the raw input is unsuccessful (**Supplementary Fig. 3a**) while the neural network recovers a clean separation between the classes (**Supplementary Fig. 3b**). In the case of image data, a deep learning model creates a nonlinear connection to a simpler latent space, where decision boundaries accurately discriminate between the complex labels. In contrast to this image-content prediction example, the annotation of cell types in single cell data relies on the cluster analysis of a feature reduced space (e.g. through selecting highly variable genes and performing PCA as a pre-processing step). Here, the cell type label is a non-complex label in the sense that the activity of a few key marker genes provides an often-sufficient grouping of similar cells, *i.e.* the distance on the reduced feature space induces an already biologically interpretable structure of the data.

Hence, to be able to use deep-learning-based representations to discover non-trivial and generalizable feature spaces, particularly interesting for subsequent transfer learning^28^, more complex tasks should be addressed. Such complex functional relationships are encountered between the cellular state (cell type or cluster identity) and functional assays, responses to drug treatment or gene knockouts^29,30^. Furthermore, complex labels such as cell fates require creative experimental methods for tracing over several generations in both cell lines and complete animals, such as traceable genetic manipulation^31,32^. In tissue samples, the transcriptome of cells depends on its direct neighbourhood^33,34^, *i.e.* when we combine spatial information with RNA-seq, we can ask to predict a cell state together as a function of its neighborhood as in local frames in image data. Furthermore, paired multi-omics data such as CITE-Seq^35^ allow the prediction of protein abundances, where deep learning models are already applied^36^. Altogether, when labels become more complex, we expect the benefits of deep learning based classifiers to increase.

While waiting for the emergence of such labels, the scientific community has developed a series of unsupervised deep-learning approaches, including batch effect correction^37^, large scale data integration^38^, single-cell perturbation prediction^39^, and dropout effects denoising^40^. However, the prediction of cell types is cell based, not cluster based. Therefore, a better clustering does not address the lack of complexity in the cell type annotation, but it reduces noise, increases accuracy and reduces occasional inconsistencies.

In general, the application of neural networks in supervised cell-type annotation provides practical advantages in terms of flexible applicability, highly optimized training libraries, and access to learned feature representation in latent spaces for downstream analysis and visualization^28^.

In the present study, we were not able to observe any significant improvement in classification accuracy based on neural networks, in contrast to other fields, such as computer vision^41^ and natural language processing^42–44^. Therefore, if we are to achieve the “ImageNet moment” in single-cell genomics, where end-to-end deep-learning models heavily outperform classical machine learning methods in the limit of large training datasets, we will need to work on better annotation and data with more covariates such as replicates, samples or patients in the future.

