## Supplementary Material for "Deep learning does not outperform classical machine learning for cell-type annotation"

### Supplementary Figures

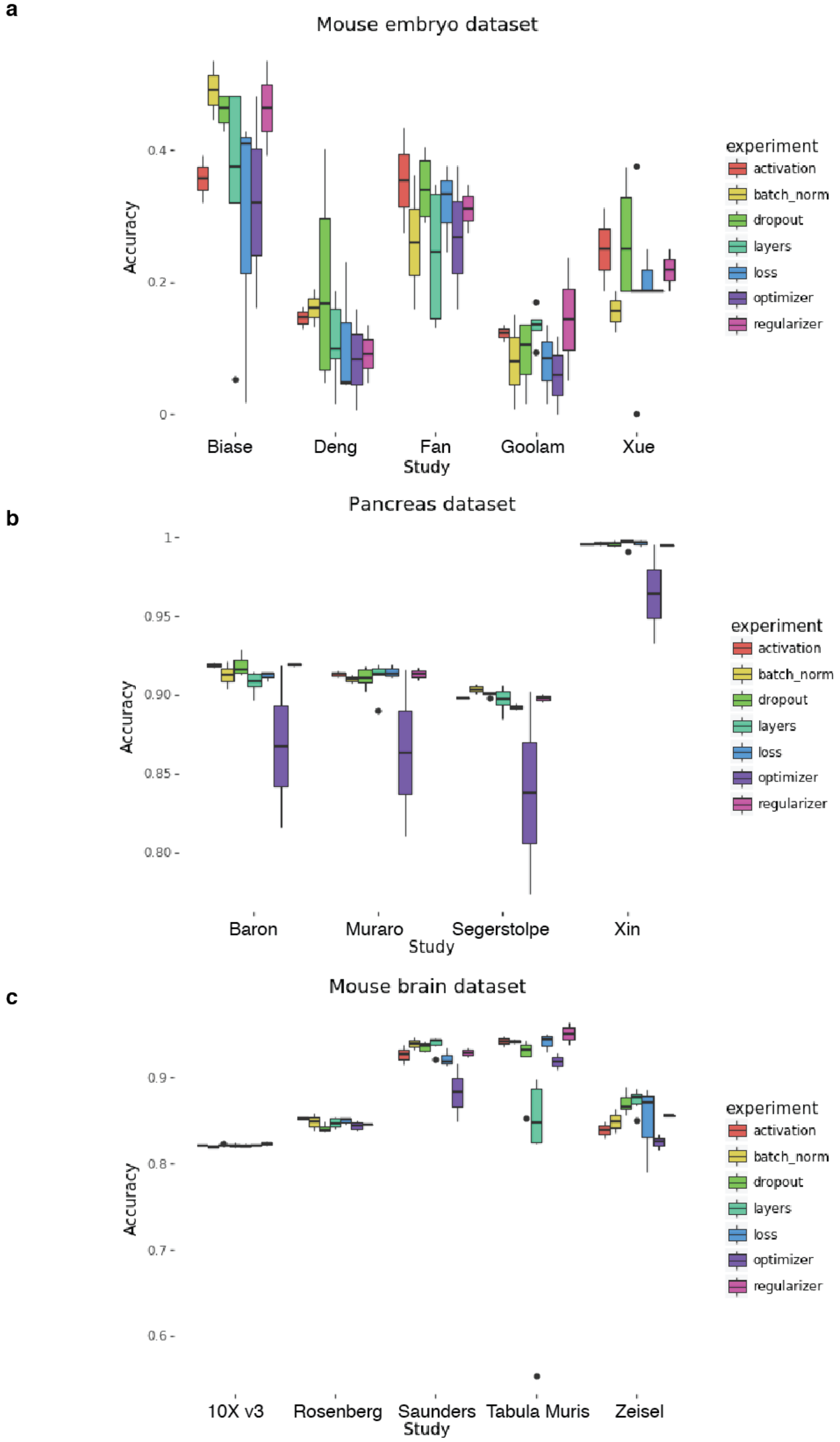


***Supplementary Figure 1: Accuracy scores for the grid search of the MLP for every data set as holdout study.*** ***(a)*** *Boxplots of accuracy scores for mouse embryonic development studies.* ***(b)*** *Boxplots of accuracy scores for human pancreas studies.* ***(c)*** *Boxplots of accuracy scores for mouse brain studies.*

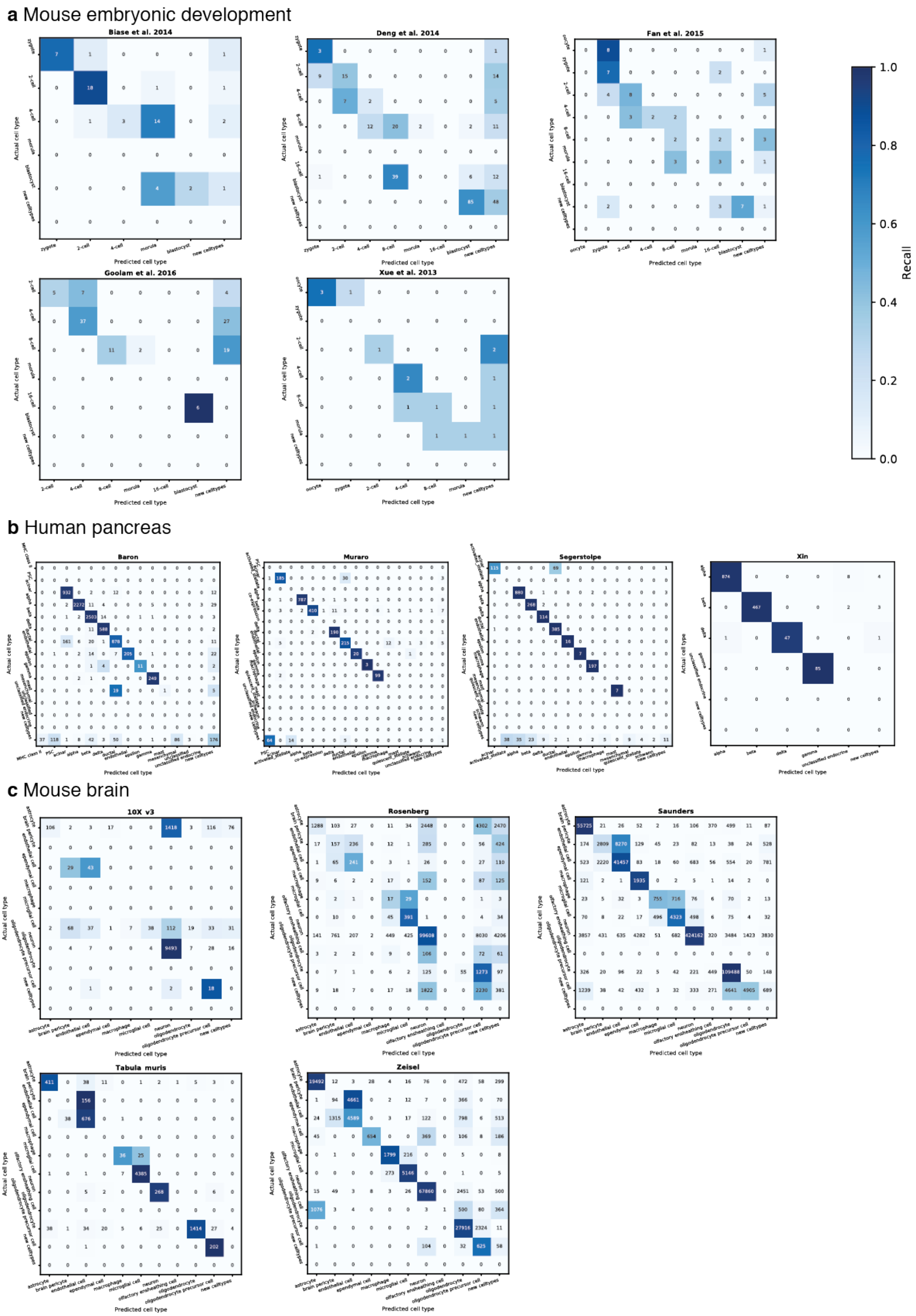


***Supplementary Figure 2: Confusion matrices for the recall of the MLP for every data set as holdout study.*** ***(a)*** *Confusion matrices for mouse embryonic development studies.* ***(b)*** *Confusion matrices for human pancreas studies.* ***(c)*** *Confusion matrices for mouse brain studies.*

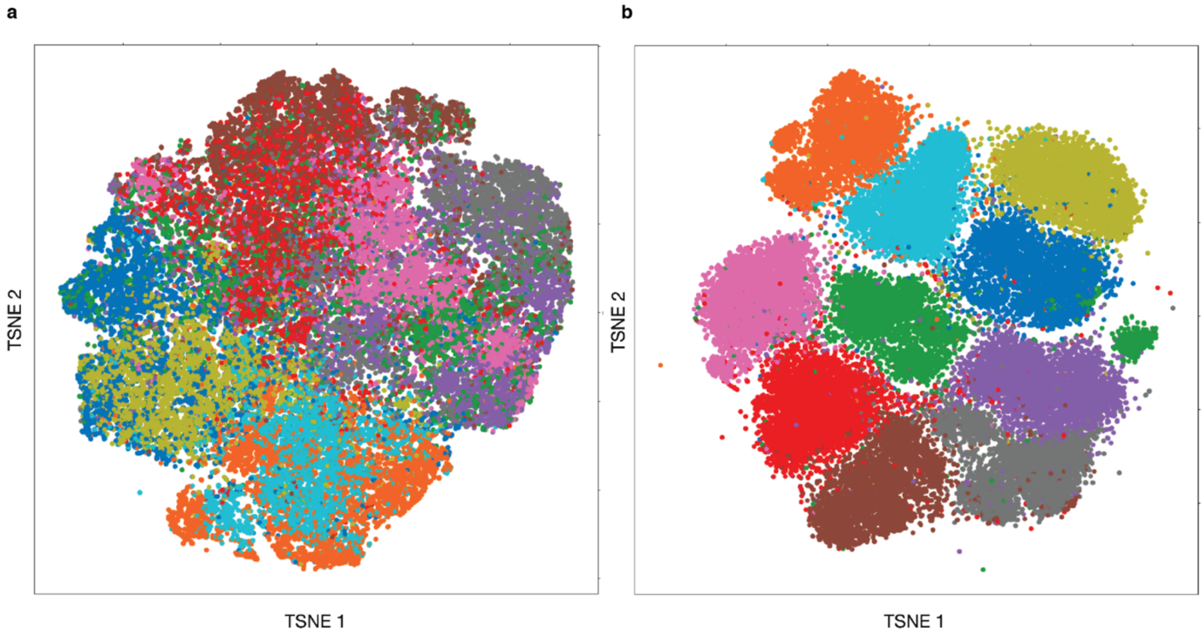


***Supplementary Figure 3: t-SNE plots of the CIFAR-10 data set, colored by the 10 classes. (a)*** *t-SNE computed on the raw input images.* ***(b)*** *t-SNE computed on the learned features of a neural network.*

### Supplementary Methods

#### Cell type annotation algorithms

Logistic regression (lgr): We used the sklearn (0.19.1) implementation for logistic regression in python 3.6.8 with the following parameters:

| Parameter | mouse embryo | human pancreas | mouse brain |
|---|---|---|---|
| C | 1 | 0.01 | 1 |
| class_weight | None | None | None |
| dual | False | False | False |
| fit_intercept | True | True | True |
| intercept_scaling | 1 | 1 | 1 |
| max_iter | 100 | 100 | 100 |
| multi_class | ovr | ovr | ovr |
| n_jobs | 1 | 1 | 1 |
| penalty | l1 | l2 | l1 |
| random_state | 42 | 42 | 42 |
| solver | liblinear | liblinear | liblinear |
| tol | 0.0001 | 0.0001 | 0.0001 |
| warm_start | False | False | False |

k-NN classifier (knn): We used the sklearn (0.19.1) implementation for the k-NN classifier in python 3.6.8 with the following parameters:

| Parameter | mouse embryo | human pancreas | mouse brain |
|---|---|---|---|
| algorithm | auto | auto | - |
| leaf_size | 30 | 30 | - |
| metric | minkowski | minkowski | - |
| metric_params | None | None | - |
| n_neighbors | 5 | 30 | - |
| p | 2 | 2 | - |
| weights | distance | distance | - |

support vector machine (svm): We used the sklearn (0.19.1) implementation for the support vector machine classifier in python 3.6.8 with the following parameters:

| **Parameter** | **mouse embryo** | **human pancreas** | **mouse brain** |
|---|---|---|---|
| C | 0.1 | 0.1 | - |
| cache_size | 200 | 200 | - |
| class_weight | None | None | - |
| coef0 | 0.0 | 0.0 | - |
| decision_function_shape | ovr | ovr | - |
| degree | 3 | 3 | - |
| gamma | auto | auto | - |
| kernel | linear | linear | - |
| probability | True | True | - |
| shrinking | True | True | - |
| tol | 0.001 | 0.001 | - |
| random_state | 42 | 42 | - |

gradient boosted trees (xgb package): We used the xgboost (0.72) implementation for the gradient boosted trees classifier in python 3.6.8 with the following parameters:

| **Parameter** | **mouse embryo** | **human pancreas** | **mouse brain** |
|---|---|---|---|
| base_score | 0.5 | 0.5 | 0.5 |
| booster | gbtree | gbtree | gbtree |
| colsample_bylevel | 1 | 1 | 1 |
| colsample_bytree | 0.75 | 0.75 | 1.0 |
| gamma | 0 | 0 | 0 |
| learning_rate | 0.1 | 0.1 | 0.1 |
| max_detla_step | 0 | 0 | 0 |
| max_depth | 4 | 4 | 4 |
| min_child_weight | 1 | 1 | 1 |

| | | | |
|---|---|---|---|
| missing | nan | nan | nan |
| n_estimators | 50 | 100 | 500 |
| reg_alpha | 0 | 0 | 0 |
| reg_lambda | 1 | 1 | 1 |
| scale_pos_weigth | 1 | 1 | 1 |
| subsample | 0.75 | 0.75 | 1.0 |
| tree_method | hist | hist | hist |

multilayer-perceptron (mlp):
We used the sklearn (0.19.1) implementation for the multilayer-perceptron classifier in python 3.6.8 with the following parameters:

| **Parameters** | **mouse embryo** | **human pancreas** | **mouse brain** |
|---|---|---|---|
| activation | elu | elu | relu |
| alpha | 0.0001 | 0.0001 | 0.0001 |
| batch_size | auto | auto | auto |
| batch_normalization | true | false | false |
| beta_1 | 0.9 | 0.9 | 0.9 |
| beta_2 | 0.999 | 0.999 | 0.999 |
| dropout_rate | 0.0 | 0.0 | 0.2 |
| early_stopping | True | True | True |
| epochs | 200 | 200 | 200 |
| epsilon | 1e-08 | 1e-08 | 1e-08 |
| hidden_layer_sizes | (100,) | (100,100) | (100,) |
| learning_rate | invscaling | invscaling | invscaling |
| learning_rate_init | 0.01 | 0.001 | 0.001 |
| loss | cos | KL | cross-entropy |
| max_iter | 200 | 200 | 200 |
| momentum | 0.9 | 0.9 | 0.9 |
| nesterovs_momentum | True | True | True |

| | | | |
|---|---|---|---|
| power_t | 0.5 | 0.5 | 0.5 |
| regularization | L1 | L2 | L2 |
| solver | adam | adam | adam |
| tol | 0.001 | 0.001 | 0.001 |
| validation_fraction | 0.1 | 0.1 | 0.1 |
| warm_start | False | False | False |

For the gridsearch model selection using an mlp with class weighting (see **Supplementary Table 1**), we used the following parameters:

| **Parameters** | **mouse embryo** | **human pancreas** | **mouse brain** |
|---|---|---|---|
| activation | relu | relu | relu |
| alpha | 0.0001 | 0.0001 | 0.0001 |
| batch_size | auto | auto | auto |
| batch_normalization | false | false | false |
| beta_1 | 0.9 | 0.9 | 0.9 |
| beta_2 | 0.999 | 0.999 | 0.999 |
| dropout_rate | 0.2 | 0.0 | 0.2 |
| early_stopping | True | True | True |
| epochs | 200 | 200 | 200 |
| epsilon | 1e-08 | 1e-08 | 1e-08 |
| hidden_layer_sizes | (100,100) | (200,200,100,100) | (100,) |
| learning_rate | invscaling | invscaling | invscaling |
| learning_rate_init | 0.01 | 0.001 | 0.001 |
| loss | KL | cross-entropy | cross-entropy |
| max_iter | 200 | 200 | 200 |
| momentum | 0.9 | 0.9 | 0.9 |
| nesterovs_momentum | True | True | True |
| power_t | 0.5 | 0.5 | 0.5 |
| regularization | L1 | L1 | L2 |

| | | | |
|---|---|---|---|
| solver | adam | adam | adam |
| tol | 0.001 | 0.001 | 0.001 |
| validation_fraction | 0.1 | 0.1 | 0.1 |
| warm_start | False | False | False |

scmap-cell: We used *scmap-cell* as implemented in scmap (v. 1.4.1) in R (v. 3.5.2) following the accompanying vignette with default parameters.

scMatch: We cloned the github repository of scMatch (version tag fbf317eca4ae964bc0cd9d9375eaa3196d72bcb8). We used jupyter notebooks to run the scMatch main function (instead of the command line tool as described in https://github.com/asrhou/scMatch). Importantly, scMatch does not learn an annotation based on n-1 reference studies, but compares a query dataset to a predefined FANTOM5 reference. The cell type annotations in our scenarios did not match well with the FANTOM5 reference in scMatch, especially embryonic tissues are only roughly resolved and did not reflect our annotation. Therefore, we modified the evaluation of the prediction with scMatch. In the embryonic dataset, we considered all predictions of embryonic tissues as correct. In the human pancreas dataset, we considered a prediction as correct, if it assigned the cell to pancreatic or endocrine tissue.

SingleR: We cloned the github repository of SingleR (version tag db4823b380ba2c3142c857c8c0695200dd1736f6). We applied a custom script for SingleR adapted from https://github.com/tabdelaal/scRNAseq_Benchmark/ [19]

Garnett: We cloned the github repository of Garnett (version tag 0.1.4). We applied a custom script for Garnett adapted from https://github.com/tabdelaal/scRNAseq_Benchmark/ [19]a marker-gene list for every cell type as input. For the embryonic development scenario, top 10 marker genes for each developmental state were derived using a t-test of each cell type vs all other cells (scanpy rank_genes_groups function). We used only genes that appeared characteristic in almost all datasets. For human pancreas, we used a custom cell type marker annotation, and in the mouse brain scenario, we simplified the mouse brain cell type markers file from the Garnett website https://cole-trapnell-lab.github.io/garnett/classifiers/, which was originally derived for the Zeisel et al. dataset. We did not use their classifier directly, because it was trained on a dataset we use in our study and it would have biased the outcome.

cellFishing.jl: We cloned the github repository of cellFishing.jl (version 0.3.0 in julia 1.4.1). We used jupyter notebooks to run cellfishing.jl as described on the github repository https://github.com/bicycle1885/CellFishing.jl . In order to assign cell types, we used a majority vote based on 10 nearest neighbors.

#### Parameter Optimization via Grid Search

We used the sklearn (0.19.1) implementation of GridSearchCV in python 3.6.8 to perform the 5-fold cross-validated Grid Search for all of our models on all three biological systems. For each classifier we defined a parameter grid.

XGBoost

| Parameter | Range |
|---|---|
| max_depth | [2,4,8,10] |
| n_estimators | [20,50,100,500] |
| subsample | [0.75,1.00] |
| colsample_bytree | [0.75,1.00] |

Logistic Regression

| Parameter | Range |
|---|---|
| C | [0.01,0.1,1,10] |
| penalty | ['l1','l2'] |

Multi-Layer-Perceptron

| Parameter | Range |
|---|---|
| hidden_layer_sizes | [500,500,450,450,450,250,250,150,150,100,100]; [200,200,100,100]; [100,100]; [100] (for mouse brain: [200,200,100,100], [100,100],[100]) |
| learning_rate_init | [0.01,0.001] |
| activation | ['relu','sigmoid', 'elu'] |
| loss | cross-entropy, cosine, KL |
| dropout rate | 0.0, 0.2, 0.4, 0.5 (for mouse brain: 0.0, 0.2, 0.5) |
| regularization | L1, L2 |
| optimizer | Adam, SGD (for mouse brain: Adam) |
| batch normalization | false, true |

Support Vector Machine

| Parameter | Range |
| --- | --- |
| C | [0.01,0.1,1,10] |
| kernel | ['poly','linear'] |

kNN-Classifier

| Parameter | Range |
| --- | --- |
| n_neighbors | [5,10,30] |
| weights | ['uniform','distance'] |

#### Data availability and pre-processing

Mouse embryonic development studies:
We used 5 different SMARTseq-based scRNA-seq studies[6–10], consisting of 56, 314, 88, 124 and 16 cells, respectively, with the following accession ids: E-GEOD-57249, E-MTAB-3321, GSE53386, E-GEOD-45719 and E-GEOD-44183. Fastq files were mapped to Ensembl[45] mouse transcriptome (version GRCm38.p5.87) with Salmon[46] (version 0.8.2, kmer = 21 to tolerate different read length). We tested the cell type annotation models on log-normalized data without further gene filtering or quality control.

Human Pancreas studies:
We used 4 different human pancreas studies[11–14], consisting of 3514 cells (Smart-Seq2), 8569 cells (inDrop), 1600 cells (SMARTer) and 2126 cells (CEL-Seq2), respectively, with the following accession ids: E-MTAB-5061, GSE84133, GSE81608 and GSE85241. Please note that a pre-processed and annotated version of the count matrices is provided by the Hemberg lab, which we used as a resource. We tested the cell type annotation models on log-normalized data without further gene filtering or quality control.

Mouse brain studies:
We used 5 different scRNA-seq studies[15–18] and a single cell gene expression dataset for 10k mouse brain cells (E18) from 10X Genomics, consisting of 133,435 cells (SPLiT-seq protocol, Rosenberg et al. (2018)), 145,954 cells (10X Genomics Chromium, Zeisel et al. (2018)), 7,856 myeloid and non-myeloid cells (SMART-Seq2 protocol, Tabula Muris (2018)), 691,489 cells (Drop-seq protocol, Saunders et al. (2018)), and 11,741 cells (10X Genomics Chromium, 10X Genomics), respectively. Overall, this scenario contains 990,475 cells.
Rosenberg et al. have deposited the raw count matrix on GEO with accession ID GSE110823. Zeisel et al. have deposited the data as web resource, from which we downloaded the annotated count matrix as loom file 'L5_all.loom'. FAC-sorted mouse brain tissue data (myeloid and non-myeloid cells) from Tabula Muris are available through figshare. Saunders et al. have deposited the data as web resource, from which we downloaded count matrices per cell type in DGE file format ('DGE by Region' section). A single cell gene expression dataset for 10k mouse brain cells (E18) was downloaded from 10X Genomics.

All datasets except for the 10X Genomics 10k mouse brain dataset were pre-filtered and annotated. We unified any differences in notation across the datasets using fuzzy string matching and log-normalized the count matrices. We used the scanpy[47] framework (v. 1.4+14.gd4a7a2d) to carry out pre-processing. We filtered the 10X Genomics 10k mouse brain dataset by removing all cells with less than 650 counts and less than 500 expressed genes. All genes detected in less than 2 counts were removed. The final count matrix was log-normalized. Then, data were clustered using louvain clustering with resolution 1.0. Further, we annotated the clusters through marker gene expression.

Cifar10 image data set:
60,000 images annotated in 10 mutually exclusive classes are available under https://www.cs.toronto.edu/~kriz/cifar.html.

#### Significance test for overall performance

We performed a paired, one-sided, pairwise wilcoxon rank sum test in R (v. 3.5.2) on all studies (n=9 for pairwise comparisons of knn, svm, lgr, xgb, mlp,scmap-cell, SingleR, Garnett, scMatch and cellFishing.jl, and n=14 for pairwise comparisons of lgr, xgb and mlp, respectively) to assess the overall performance of the classifiers with Benjamini-Hochberg correction for multiple testing (function pairwise.wilcox.test).

#### Supplementary Tables

**Supplementary Table 1:** Impact of class weighting on accuracy in the multilayer perceptron compared to model selection based on ROC-AUC per cell type. Using ROC-AUC improved overall accuracy and average accuracy in almost all cases.

| | Class weighting MLP | | ROC-AUC selected MLP | |
|---|---|---|---|---|
| Dataset | Accuracy (overall) | Average Accuracy | Accuracy (overall) | Average Accuracy |
| Biase | 0.714 | 0.631 | 0.536 | 0.528 |
| Deng | 0.327 | 0.352 | 0.425 | 0.392 |
| Fan | 0.246 | 0.204 | 0.377 | 0.337 |
| Goolam | 0.144 | 0.145 | 0.449 | 0.309 |
| Xue | 0.250 | 0.250 | 0.500 | 0.483 |
| Baron | 0.894 | 0.795 | 0.917 | 0.797 |
| Muraro | 0.895 | 0.980 | 0.898 | 0.940 |
| Segerstolpe | 0.921 | 0.974 | 0.987 | 0.984 |
| Xin | 0.991 | 0.994 | 0.991 | 0.994 |
| 10X Genomics (v3) | 0.822 | 0.423 | 0.826 | 0.524 |
| Rosenberg | 0.717 | 0.415 | 0.779 | 0.330 |
| Saunders | 0.935 | 0.716 | 0.934 | 0.732 |
| Tabula Muris | 0.949 | 0.772 | 0.941 | 0.781 |
| Zeisel | 0.736 | 0.612 | 0.878 | 0.655 |

**Supplementary Table 2:** Benjamini-Hochberg corrected p-values of the paired, one-sided, pairwise wilcoxon rank sum test with the alternative hypothesis “Method in row performed with less accuracy than method in column”. Data: Mouse embryonic development and human pancreas studies (n=9). Significant p-values (p<0.01) are highlighted in boldface.

| | knn | lgr | svm | xgb | mlp | scmap-cell | SingleR | Garnett | scMatch |
|---|---|---|---|---|---|---|---|---|---|
| lgr | 0.9321 | | | | | | | | |
| svm | 0.9321 | 0.2539 | | | | | | | |
| xgb | 0.7910 | 0.6252 | 0.7584 | | | | | | |
| mlp | 0.3516 | 0.0732 | 0.2539 | 0.1934 | | | | | |
| scmap-cell | **0.0098** | **0.0098** | 0.0292 | **0.0098** | 0.1934 | | | | |
| SingleR | 0.3516 | 0.1285 | 0.4583 | 0.2539 | 0.7584 | 0.9791 | | | |
| Garnett | **0.0098** | **0.0098** | **0.0098** | **0.0098** | **0.0098** | 0.0176 | **0.0098** | | |
| scMatch | 0.3516 | 0.3685 | 0.3516 | 0.3516 | 0.3685 | 0.4134 | 0.3685 | 0.9791 | |
| cellFishing.jl | 0.7031 | 0.6606 | 0.7584 | 0.6252 | 0.9791 | 1.0000 | 0.9791 | 1.0000 | 0.9465 |

**Supplementary Table 3:** Benjamini-Hochberg corrected p-values of the paired, one-sided, pairwise wilcoxon rank sum test with the alternative hypothesis “Method in row performed with greater accuracy than method in column”. Data: Mouse embryonic development and human pancreas studies (n=9). Significant p-values (p<0.01) are highlighted in boldface.

| | knn | lgr | svm | xgb | mlp | scmap-cell | SingleR | Garnett | scMatch |
|---|---|---|---|---|---|---|---|---|---|
| lgr | 1 | | | | | | | | |
| svm | 1 | 1 | | | | | | | |
| xgb | 1 | 1 | 1 | | | | | | |
| mlp | 1 | 1 | 1 | 1 | | | | | |
| scmap-cell | 1 | 1 | 1 | 1 | 1 | | | | |
| SingleR | 1 | 1 | 1 | 1 | 1 | 1 | | | |
| Garnett | 1 | 1 | 1 | 1 | 1 | 1 | 1 | | |
| scMatch | 1 | 1 | 1 | 1 | 1 | 1 | 1 | 1 | |
| cellFishing.jl | 1 | 1 | 1 | 1 | 0.923 | 0.615 | 0.923 | **0.088** | 1 |

**Supplementary Table 4:** p-values of the paired, one-sided, pairwise wilcoxon rank sum test with the alternative hypothesis “Method in row performed with less accuracy than method in column”. Data: Mouse embryonic development, human pancreas studies and mouse brain studies (n=14).

| | lgr | mlp |
|---|---|---|
| mlp | 0.31 | |
| xgb | 0.31 | 0.31 |
